# Convergence on BRAF and MAPK Signaling in Glioma Development in a P53-ENU model

**DOI:** 10.64898/2026.02.23.707201

**Authors:** Kinjal Desai, Anna Tymofyeyeva, Margaret Javier, Nataliia Svergun, Vincent Ye, Garrett Bullivant, Jannine Forst, Matthaeus Ware, Michelle Kushida, Lilian Lee, Chunying Yu, Heather Whetstone, Ryan Ward, Biren M. Dave, Ahmed Aman, Peter B. Dirks

## Abstract

Pediatric high-grade gliomas (HGGs) are aggressive and lethal brain tumors that account for 15– 20% of all pediatric central nervous system (CNS) malignancies and remain largely incurable. These tumors, despite having mutational targets that activate MAPK signaling, are frequently resistant to targeted therapies in their malignant states, but often show responses when they are in lower grade form. These findings suggest a need to identify and intercept early tumorigenic events that arise in earlier tumor developmental stages. To investigate the molecular events in glioma progression, we developed and characterized a *Nestin^Cre/+^*;*Trp53^fl/fl^* mouse model combined with *in utero* exposure to the classic chemical mutagen *N*-ethyl-*N*-nitrosourea (ENU). This model mimics the context of genetic predisposition paired with an environmental genotoxic insult, and mice with loss of TRP53 in early neural precursors who are exposed to ENU at embryonic day 13.5 reproducibly develop HGGs postnatally that retain features of the human tumors. By sampling discrete lesions at premalignant, early, and late stages, we observed progressive increases in genetic complexity, stemness features, and immune signatures across tumor evolution. Notably, a recurrent *Braf* mutation emerged in a majority of early lesions and persisted in advanced tumors, consistent with the occurrence of BRAFV600E mutations in human gliomas that arise in children and undergo transition from lower grade to higher grade stages. Additional components of the RAS-RAF-MAPK signaling cascade, including *Kras*, *Nras* and *Nf1* were found to be mutated at the late tumor stage, indicating convergent activation of this pathway in this model. Cell lines derived from early lesions responded to BRAF inhibitors, but cells from endpoint tumors were less responsive. Together, this model reveals aspects of the molecular and cellular evolution of glioma development *in vivo*, and identifies RAS-MAPK signaling as a critical molecular bottleneck selected for the ENU-induced mutations. This genetic-environmental model may be valuable to understanding key determinants of glioma initiation and progression, and for evaluating new therapies that limit MAPK signaling.

## Introduction

Pediatric gliomas constitute a diverse spectrum of neoplasms and represent common malignant brain tumors in children, particularly when they occur in the cerebral hemispheres and brainstem [1,2]. Among these, pediatric high-grade gliomas (pHGGs) are highly aggressive malignancies, accounting for roughly one third of cases, and representing a leading cause of cancer-related mortality in children and young adults [3,4]. Despite significant advances in cancer research and clinical care over the past four decades, which have significantly increased survival rates for other childhood cancers, pHGG survival has remained virtually unchanged [5]. These tumors tend to develop resistance even to targeted treatments at late stages [6], underscoring a critical need for treatments at early stages of disease when they are fueled by fewer driver mutations and less cellular heterogeneity.

Although pHGGs share morphological similarities with adult HGGs, their genetic and molecular landscapes have unique features [18,19,20,21]. pHGGs may be further sub-classified into diffuse midline gliomas, which are characterized by H3K27 alterations, hemispheric HGGs that often harbor H3G34 mutations, and diffuse pediatric-type high-grade glioma, H3-wildtype and IDH-wildtype [22,23,22]. Compared to adult HGG, EGFR amplification and PTEN loss/mutation are less common [24] and TERT promoter mutations are rare [25] in pHGGs. Copy number changes are not as frequent in pediatric disease; less frequent gains of chromosome 1q and losses of chromosomes 13q and 14q are seen in pHGG, rather than chromosome 7 gain and chromosome 10 loss observed in adult GBM [26]. Molecular profiling over the last 15 years have also identified oncogenes frequently found in childhood onset lower grade gliomas. Apart from *TP53*, these include *PDGFRA* [27], *ATRX* [26], and *BRAF* [9,28,29]. Notably, BRAF mutations represent a molecularly distinct subgroup that has important implications for tumor biology and progression. *BRAFV600E* mutations are present in 15–20% of pediatric low-grade gliomas [30,31] and 5–10% of pHGG [26,32,33]. *BRAF* is a key component of the RAS/RAF/MAPK pathway, which transmits extracellular signals from the cell membrane to the nucleus. Crosstalk between this pathway and other critical signaling cascades regulates cell growth, differentiation, and survival. Activation of *BRAF* by point mutation is a common driver of transformation from pediatric low-grade glioma to pediatric high-grade glioma (pHGG) [31,34,35], with or without cooperating loss of CDKN2A.

Genetically engineered mouse models (GEMMs) are highly tractable systems to study tumor development from the earliest stages. GEMMs offer several advantages over xenograft or culture models, including an intact immune system and a preserved blood-brain barrier, capturing endogenous microenvironmental cues and intrinsic developmental contexts that shape tumor initiation and evolution [7,8]. Currently, most existing glioma GEMMs rely on combining multiple simultaneous genetic lesions to trigger neoplastic transformation. Common strategies include *Trp53* inactivation with *Nf1* and *Pten* loss [9,10,11], *Trp53* deficiency with *Atrx* loss [12], *Egfr* mutations with *Cdkn2a* loss [13], and *Pdgfra/b* overexpression with *Trp53* or *Cdkn2a* co-deletions [14]. Although these models are extremely valuable, they undoubtedly fall short of modeling early steps of disease, which are thought to involve sequential activation of genetic changes and subsequent microenvironmental selection. Conditional *Trp53* deletion has been a particular focus, given its central role as a tumor suppressor and the fact that over a third of human pHGGs exhibit evidence of TP53 inactivation [15], and that patients with germline TP53 alterations, such as in Li Fraumeni Syndrome, are susceptible to glioma that inevitably becomes malignant. In our own lineage-tracing mouse studies, coupled with genomic and longitudinal studies of human tumors, there is evidence to support that HGG arise clonally and over a relative long timeframe, acquiring mutations progressively [16,17].

The mutagen *N*-ethyl-*N*-nitrosourea (ENU) is a potent alkylating agent that introduces point mutations by ethylation of DNA bases, with a tissue-specific preference for GC→AT transitions [36,37,38]. Beyond mutagenesis, ENU may effect additional mechanisms such as increasing cellular oxidative stress, and suppressing the bone marrow and immune system to contribute to malignant disease [39]. Epidemiological data has determined that maternal dietary intake of *N*-nitroso compounds and precursors of *N*-nitroso compounds, including cured meats and oil products, during pregnancy increases the risk of brain tumors in offspring [40]. The use of ENU to induce brain tumors has been reported since the late 1960s, with single injections to pregnant rats, transplacental administration, or administration during the first week of birth resulting in a high incidence of gliomas [36,41,42]. In mouse models, however, administration of ENU alone is insufficient to produce gliomas and requires additional genetic manipulation, such as loss of function of *Trp53* [43].

Here, we focused on tumor initiation in Nestin^+^ cells through development. *Nestin* is expressed in neural stem and progenitor cells early in brain development and persists in stem cell niches postnatally [44]. Using the Cre-loxP system, we floxed out *Trp53* in *Nestin*-expressing neuronal precursor cells, which are also considered candidate cells of origin for pHGG [45,46,47,48].

Administration of ENU at embryonic day 13.5 (E13.5) results in 100% brain tumor incidence in the mice by six months of age, with 90% of animals having a single tumor. By serial MRI imaging we saw that mice typically had a single lesion at 12–15 weeks that slowly grew until very rapid tumor growth occurred over a short period, at which point animals needed to be euthanized. Whole exome sequencing (WES) and RNA sequencing at different stages of disease confirmed alterations in key genetic pathways characteristic of pHGG, particularly with a large proportion of gain-of-function mutations in the RAS/RAF/MAPK pathway in early lesions.

Models of gliomagenesis need to take into consideration both underlying genetic changes and microenvironmental influences that drive pre-neoplastic cells into fully malignant cells. By exposing *Trp53*-mutant embryonic neural stem cells to a single dose of ENU, an alkylating agent but also representative of an environmental mutagen, we generated a reliable model of glioma that progresses through the stages of tumor progression: premalignancy to malignancy. The mutations in MAPK pathway that are seen in a high proportion of tumors reinforce that activation of this signaling pathway is a critical feature, and maybe even a necessary bottleneck, for gliomagenesis. Our model, which generates variable disease states within a defined limit, could prove very useful for devising therapies that target full blown tumors as well as their earlier, and simpler forms. This model may allow for therapies that intercept disease and maybe even prevent full transformation.

## Materials and methods

### Mouse genetics and housing

All mouse experiments were approved by the Hospital for Sick Children’s Animal Care Committee and followed all legal and ethical regulations. Transgenic *Nestin*-Cre mice were purchased from the Jackson Laboratory (JAX #003771). *Trp53^fl/fl^* mice were provided by Dr. Chi-Chung Hui, Toronto, Hospital for Sick Children (SickKids). Mice were bred in-house at the Laboratory Animal Services (LAS) Facility, SickKids, and at three weeks of age, litters of mice had their ears notched for subsequent genotyping as recommended by the Jackson Laboratory, except *Trp53^fl/fl^* mice, which were genotyped based on Chow *et al*., 2011.

At 6 weeks of age, the *Nestin^Cre/+^*;*Trp53^fl/fl^*; ENU^+^ (NCpE) mice were transferred from the animal facility at the Hospital for Sick Children to the Spatio-Temporal Targeting and Amplification of Radiation Response (STTARR) Facility, University Health Network, Toronto. At both facilities, the mice had free access to rodent chow and water. They were housed in a 14-hour light, 10-hour dark cycle room with ambient temperature at 22–24 °C and 45–50% humidity (LAS); and in a 12-hour light, 12-hour dark cycle room with ambient temperature at 21–22 °C and 45–60% humidity (STTARR). The mutant mice that were expected to develop tumors were monitored daily and euthanized once they developed end-stage symptoms (i.e., domed head, dehydration, ataxia). Our study included both male and female mice, with the effect of sex not being evaluated as a specific biological variable.

### Timed breeding and ENU administration

After 5 weeks of age, *Trp53^fl/fl^* females were used for timed mating with *Nestin^Cre/+^*;*Trp53^fl/fl^*males. One to two females were placed in a cage with one male in the evening (after 17:00) for mating. The following morning, the females were checked for copulatory plugs, weighed, and separated from the male. The females were weighed regularly until pregnancy was confirmed. Pregnant females at embryonic day 13.5 (E13.5; morning of 13 days after plug detection) received a single intraperitoneal injection of freshly prepared *N*-ethyl-*N*-nitrosourea (ENU; Sigma-Aldrich; 25 mg/kg) in saline.

### Histological characterization

#### Tissue Processing

Tissue from postnatal mice was dissected after transcardial perfusion of PBS followed by 4% PFA, and subsequently fixed from 24–72 hours at room temperature in 4% PFA or 10% formalin. The tissue was then dehydrated in ethanol, wax embedded, and cut coronally at a thickness of 5 μm on a microtome.

#### Immunohistochemistry and histology

FFPE sections were de-waxed in xylene and rehydrated in graded EtOH solutions (100%, 95%, and 75%). Antigen retrieval was then achieved via heat-induced epitope retrieval (HIER). This was performed by boiling sodium citrate buffer (10 mM Sodium citrate, 0.5% Tween-20, pH 6.0) under pressure in a microwave for 5 minutes. Bloxall (Vector Laboratories, VECTSP6000) was used for 15 minutes at room temperature to block endogenous peroxidase activity before tissues were incubated with primary antibodies overnight. A polymer-based, enzymatic secondary system was utilized to amplify antigen detection and reduce background. For primary antibodies raised in rabbit, rat, and mouse, the following ImmPRESS-HRP (Peroxidase) Polymer Detection Kits (Vector Laboratories) were used: Goat anti-rabbit (MP-7451-50), Goat anti-rat (MP-7444-15), and Horse anti-mouse (MP-7402-50). For antibodies raised in goat, a separate avidin-biotin complex (ABC) system using Horse anti-goat IgG antibody (H+L), biotinylated (Vector Laboratories, BA-9500) and the VECTASTAIN Elite ABC-HRP Reagent (Peroxidase) (Vector Laboratories, PK-7100) was utilized to achieve the same. The staining was visualized with a DAB Substrate Kit, Peroxidase (HRP) (Vector Laboratories, SK-4100) and counterstained with Mayer’s Hematoxylin (Electron Microscopy Sciences, 26043-06). For all experiment sections, PBS was used as a control. Hematoxylin and Eosin (H&E) staining for histology was performed using standard methods. In all cases, analyses were performed on three biological replicates. H&E and IHC Images were acquired using the Nikon Eclipse Ci light microscope with a Nikon DS-Fi2 camera.

The following antibodies and dilutions were used for immunohistochemistry staining: Gt anti-Sox2 (1:1000, R&D Biosystems, AF2018), Rb anti-GFAP (1:100, Dako, Z0334), Rt anti-Ki67 (1:750, Invitrogen, 740008T), Rb anti-Olig2 (1:200, Millipore Sigma, AB9610), Ms anti-CD3 (1:100, Dako, M7254), Rb anti-Iba1 (1:4000, Abcam, ab178847).

### Magnetic resonance imaging

All MR imaging was performed at the Spatio-Temporal Targeting and Amplification of Radiation Response (STTARR; www.sttarr.ca) facility, University Health Network, Toronto as previously reported. [A 7 Tesla preclinical system was used (Biospec 70/30 USR, Bruker Corporation, Ettlingen, DE), equipped with the B-GA12 gradient coil insert, 7.2 cm inner diameter RF transmit coil, plus dedicated murine brain RF receiver coil and its associated slider bed. Mice were anaesthetised and maintained at 1.8% isoflurane throughout experimentation, on the provided slider bed. Respiration was monitored via a pneumatic pillow taped above the mouse (Model 1030 monitoring and gating system, SA Instruments, Stony Brook, NY). Tumors and brain anatomy were visualised via 2D-T2-weighted imaging in the horizontal plane of the mouse brain. Data processing and visualisation were performed using the National Institutes of Health MIPAV (Medical Image Processing, Analysis, and Visualization) application. Image quantification was performed by measuring # of voxels of the identified lesion regions by contour and slice using the MIPAV statistics generator function.

### Tumor processing

Tumor or lesion locations were first identified using MRI scans obtained at the STTARR facility, and analyzed in the MIPAV tool. Distance coordinates (anterior–posterior, right–left, superior– inferior) were obtained compared to anatomical landmarks (e.g., olfactory bulbs, midline, cerebellum) to determine precise localization. The ruler tool in MIPAV was used to confirm measurements. Mice were euthanized by transcardial perfusion, and brains were dissected and placed in ice-cold PBS for transport to The Hospital for Sick Children. Brains were sectioned into 1 mm coronal slices on ice and examined under a dissecting microscope (Leica, MZ7.5).

Lesions were isolated using a 1 mm biopsy punch, while larger tumors were dissected using sterile razor blades or microscissors. Tissue samples were kept on ice and transferred to a biosafety cabinet for dissociation. Samples were incubated in Accutase (STEMCELL Technologies, 7920) at 37 °C for enzymatic dissociation, followed by gentle mechanical dissociation with a pipette. The suspension was diluted with DMEM, centrifuged at 1000 rpm for 5 min, and the supernatant was discarded. The pellet was resuspended in neurosphere (NS) media consisting of Neurocult NS-A Basal media (STEMCELL Technologies, 05700) supplemented with 2 mmol/L L-Glutamine, N2, B27, 75 µg/ml bovine serum albumin, 10 ng/ml recombinant human EGF (rhEGF), 10ng/ml basic fibroblast growth factor (bFGF and 2 µg/ml heparin.Viable cells were counted using a Countess Automated Cell Counter (Thermo Fisher Scientific). Cells were plated on poly-L-ornithine/laminin–coated 6 cm or 10 cm dishes (1:100 dilution of PSF). Cultures were maintained at 37°C and 5% CO₂, fed every 2–3 days, and passaged upon reaching confluence.

### Patient sample-derived cell lines

Fresh tumor samples were obtained from patients during their operative procedure following informed consent. All experimental procedures were performed in accordance with the Research Ethics Board at The Hospital for Sick Children (Toronto, Canada) and University of Toronto (Toronto, Canada). GSCs and HF samples were derived as previously described [49,50].

Each sample ID follows the structure GXXX, or HFXXX, where XXX is the numerical identifier of the primary sample. G: Glioblastoma neural stem cell lines grown adherently; HF: Human fetal neural stem (HFNS) cell lines.

GXXX and HFXXX samples were grown adherently in serum-free medium as described previously [49]. Briefly, cells were grown on PRIMARIA^TM^ culture plates (Corning) coated with poly-L-ornithine (Sigma) and laminin (Sigma) and maintained in Neurocult NS-A basal medium (human) (StemCell Technologies) containing 2 mM L-glutamine (Wisent), 75 µg/mL bovine serum albumin (Life Technologies), in-house hormone mix equivalent to N2 (home-made), B27 supplement (Life Technologies), 10 ng/mL recombinant human epidermal growth factor (rhEGF; Sigma), 10 ng/mL basic fibroblast growth factor (bFGF; StemCell Technologies), and 2 µg/mL heparin (Sigma). Cells were passaged using enzymatic dissociation with Accutase (StemCell Technologies).

### Inhibitor assays

The three BRAF inhibitors (Naporafenib, Exarafenib, Belverafenib) were obtained from the Drug Discovery Program, Ontario Institute for Cancer Research (OICR), Toronto. The MEK inhibitors were obtained from Cedarlane/MedChemExpress (Cobimetinib, HY-13064; Tremetinib, HY-10999). 1 mg of each inhibitor was resuspended in DMSO and stored at −20°C. Short-term working stocks of 1 mM in 10 µL were aliquoted and used for the assay. Flat bottom 96-well plates (Corning Falcon, 353072) were coated at least one day before seeding cells.

Mouse-derived glioma stem cells were seeded at 500 cells/well in 50 µL of NS media (STEMCELL Technologies, 05700); human-derived glioma stem cells previously reported in Richards et al. 2021 [51] were seeded at 1500 cells/well in 50 µL of NS media (STEMCELL Technologies, 05750). Cells were treated with each inhibitor in triplicate at final concentrations of 1 µM or 5 µM, with DMSO serving as the vehicle control. Plates were incubated for 10 days, after which alamarBlue reagent (Invitrogen, DAL1100) was added to each well (10 µL per 100 µL total volume). After 6 hours of incubation at 37 °C, fluorescence was measured using a Varioscan LUX microplate reader (Thermo Fisher Scientific). Fluorescence values were normalized to the mean of the DMSO control for each cell line, and relative cell viability was plotted using R version 4.4.3.

### RNA sequencing

Tumor samples were processed as detailed in “tumor processing.” RNA was extracted using the RNeasy Plus Mini Kit (Qiagen Cat# 74134) with on-column DNase digestion using the gDNA Eliminator Columns. RNAseq libraries were constructed using the NEB Ultra II Directional polyA mRNA Library Prep Kit (NEB Cat# E7765), following the manufacturer’s instructions.

RNA was sequenced using the NovaSeq 6000 System (Illumina) on the SP flow cell. Sequencing was done as paired-end reads with a read length of 100 bases. Raw sequencing information was collected in the form of fastq files, generated using bcl2fastq2 (v2.20). The fastq files were then assessed with FastQC (v0.12.1) [52] and MultiQC (v1.25.1) [53]. The fastqs were mapped to a decoy-aware GRCm39 transcriptome using Salmon (v1.10.0) [54] before transcript quantification by tximeta (1.24.0) [54,55], annotation retrieval by biomaRt (v2.62.1) [56], and differential expression analysis using DESeq2 (v1.46.0) [57]. Plots were created using pheatmap (v1.0.13)[58] and ggplot2 (v4.0.0) [59].

### Whole exome sequencing

Tumor samples were processed as detailed in “tumor processing.” For premalignant samples, a half hemisphere was dissected. Sequencing was performed by The Centre for Applied Genomics (TCAG). Genomic DNA samples were quantified using a Qubit High Sensitivity Assay. 500 ng of DNA was used as input material for library preparation using the Agilent SureSelect XT Reagent kit. In brief, 500 ng of genomic DNA was fragmented to 200 bp on average using a Covaris LE220 instrument. Sheared DNA was end-repaired and 3’ ends adenylated prior to ligation of adapters with overhang-T. The resulting DNA library was amplified by PCR using 10 cycles and hybridized with biotinylated probes that target mouse exonic regions. Enriched exome libraries were then amplified with 8 bp barcoded primers using an additional 10 cycles of PCR. Final libraries were validated on a Bioanalyzer 2100 DNA High Sensitivity chip (Agilent Technologies) to check for size, and quantified by qPCR using the Kapa Library Quantification Illumina/ABI Prism Kit protocol (KAPA Biosystems). Libraries were pooled in equimolar quantities and paired-end sequenced on the S4 flowcell with the V1.5 sequencing chemistry on an Illumina NovaSeq 6000 platform following Illumina’s recommended protocol to generate paired-end reads of 150 bases in length.

WES analysis was completed by the Centre for Applied Genomics (TCAG) at the Hospital for Sick Children. First, bcl2fastq2 (vv2.20) was used to demultiplex the fastqs. Following GATK best practices, the reads were aligned to the GRCm38 reference genome using BWA-MEM (0.7.12) [60]. Duplicate reads were marked using Picard Tools (v2.5.0)[61], after which GATK (v3.7) [62] was used for local realignment and base quality score recalibration, followed by variant calling using GATK HaplotypeCaller on the Sanger Mouse Genome Project V5 dataset. Finally, variants were hard filtered using the standard GATK hard filtering cutoffs. Annotation was completed using a custom Annovar-based annotation pipeline from TCAG (v27.3) that also implements VEP (release 102) and Manta (v1.6.0) [63].

### Copy number analysis

Copy number analysis was performed using CNVkit. A BED file of baited regions was prepared using the genomic regions from the Exome sequencing capture kit (Agilent). For the reference genome, we used the same mm10 that was used in exome sequence alignment. We combined normal samples that were used in our PoN into a pooled reference. The default threshold method was used to apply a fixed log2 ratio cutoff value for each integer copy number state. The “heatmap” command was used to obtain a per segment overview of large-scale CNAs across experimental cohorts, de-emphasizing low-amplitude segments. The “genemetrics” function was used to identify targeted genes with copy number gain or loss above or below the default threshold.

## Results

### Development of a novel mouse model of gliomagenesis

To model spontaneous development of gliomagenesis in an immunocompetent context, we combined neural stem cell compartment-specific deletion of *Trp53* (*Nestin^Cre/+^*;*Trp53^fl/fl^*) with *in utero* exposure to the alkylating mutagen N-ethyl-N-nitrosourea (ENU) (Fig. 1A). While *Trp53*-deleted mice alone developed brain tumors with incomplete penetrance and long latency (median survival: 321 days), ENU-exposed *Nestin^Cre/+^*;*Trp53^fl/fl^*(hereafter referred to as NCpE) mice exhibited fully penetrant glioma formation with significantly reduced survival (median survival 119.5 days; *p* < 0.0001) (Fig. 1B; Supplementary Fig. 1A–B). No difference was noted between female and male survival in this model (Supplementary Fig. 1C).

**Figure 1.**
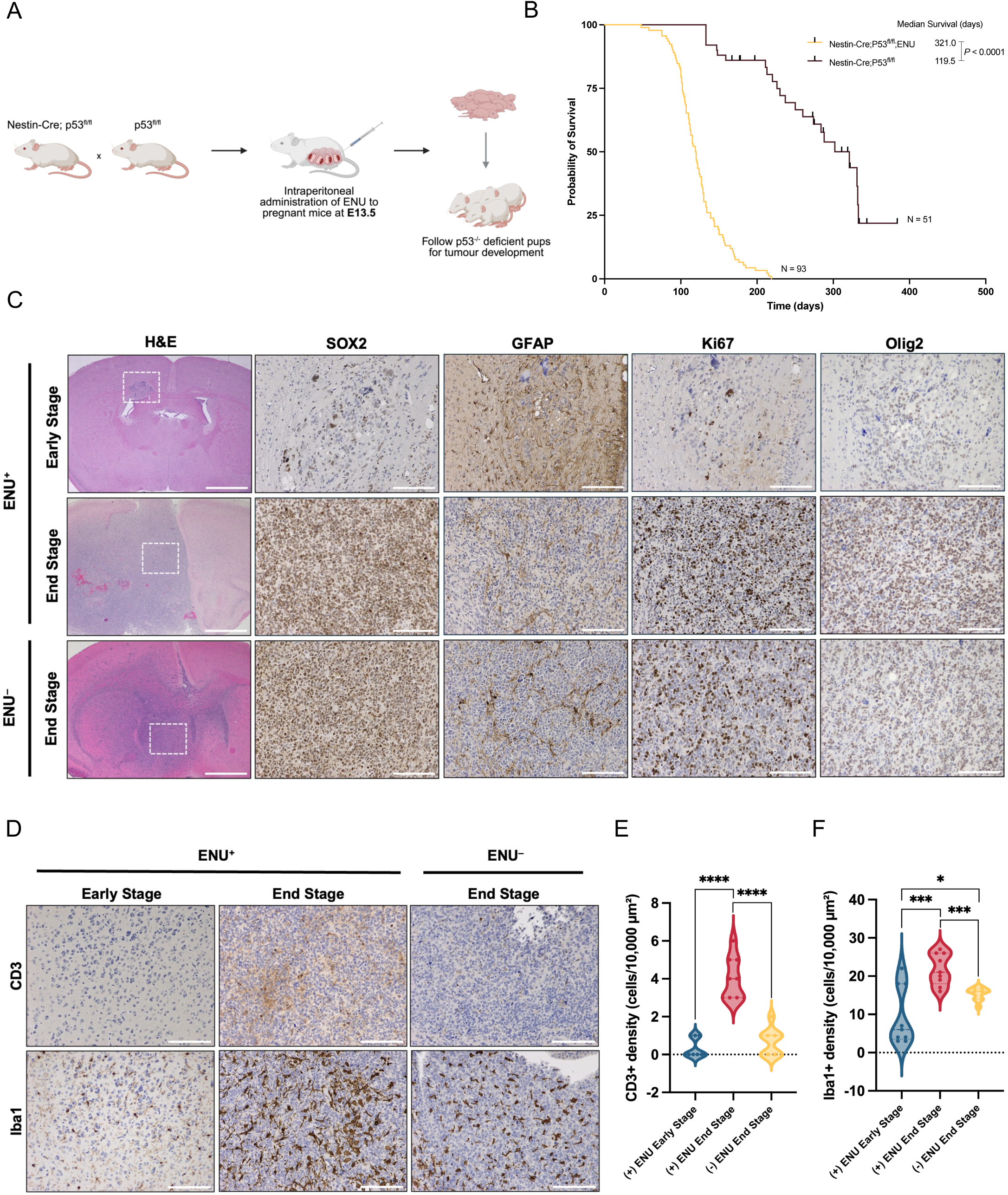
Generation of a *Nestin^Cre/+^*;*Trp53^fl/fl^*; ENU model to recapitulate immune-competent gliomagenesis. (**A**) Schematic of model generation. Pregnant *Nestin^Cre/+^*;*Trp53^fl/fl^*(NCp) dams are intraperitoneally injected with ENU at embryonic day 13.5 (E13.5), and their offspring are monitored for spontaneous tumor formation. (**B**) Kaplan-Meier survival curve of NCp with and without ENU (25 mg/kg) exposure (NCp: n=51, NCpE: n=93). Significance was estimated using the log-rank (Mantel-Cox) test. Chi square =108.7, *p*<0.0001. (**C**) Whole brain tissue sections collected at early and end stages from NCp and NCpE mice stained for H&E, Sox2, GFAP, Ki67, and Olig2. Dotted white boxes on H&E images represent the region in which the Sox2, GFAP, Ki67, and Olig2 images were acquired. H&E scale bars: 1000 µm, all other scale bars: 100 µm. (**D**) Representative immunohistochemistry (IHC) images of CD3 and Iba1expression in early or late stage NCp and NCpE mice to characterize the immune microenvironment. Scale bars: 100 µm. (**E-F**) Quantification of CD3+ T-cells (**E**) and Iba1+ microglia/macrophages (**F**) present in the early-stage lesion or end-stage tumor core. For the quantification, three 100 µm x 100 µm regions were evaluated from three independent biological samples per group. Dotted lines within the violin plots denote the median. Two-tailed unpaired *t-test*; ns *p*≥0.05, \**p*=0.0377, \*\**p*<0.0006, \*\*\*\**p*<0.0001.

We profiled early-stage lesions and end-stage tumors from mice, and generated primary cell cultures from both stages of tumor development. The presence of early-stage lesions and their locations were first detected using magnetic resonance imaging (MRI) scans (0.10–5.0 mm^3^; weeks 12–15), and end-stage was determined by the onset of neurological symptoms and confirmed using MRI (average volume: 69.780 mm^3^; week 20+). Samples were dissected at either time point for downstream analysis. A histopathological assessment of the tumors exhibited hallmark features of human high-grade gliomas, including hypercellularity, nuclear atypia, necrosis, microvascular proliferation, and a high mitotic index (Fig. 1C; Supplementary Fig. 1D). Immunohistochemistry confirmed strong expression of stemness (*Sox2*), mitotic (*Mki67*), proliferating progenitor/glioma (*Olig2*) and glial (*Gfap*) markers, with end-stage ENU^+^ (NCpE) tumors showing significantly higher expression of *Sox2*, *Olig2*, and *Mki67*, and lower expression of *Gfap* compared to both ENU^−^ (NCp) tumors and early-stage samples (Fig. 1C, Supplementary Fig. 1E), indicating an increase in stem-like and proliferating cells in end-stage tumors, specifically those induced by ENU.

### NCpE tumors have an enhanced immune signature compared with NCp tumors

We probed for the presence of T cells (CD3) and tumor-associated microglia/macrophages (*Iba1*) by performing immunofluorescence staining with specific antibodies. We observed both CD3^+^ and Iba1^+^ positive cells in early lesions, indicating activation of lymphocyte and microglial-macrophage responses to small lesions. We found that NCpE endpoint tumours had relatively low numbers of CD3^+^ cells (<5%), consistent with previous reports of low T-cell infiltration in pHGGs compared to other brain tumor types, including pLGGs or diffuse midline gliomas [14,64] (Fig. 1D,E). In contrast, Iba1^+^ cells were more abundant (∼20%) in endpoint NCpE tumors, indicating high tumor-associated microglial-macrophage infiltration (Fig. 1D,F). Furthermore, CD3^+^ as well as microglial-macrophage cells were also significantly higher in ENU^+^ compared ENU^−^ end-stage tumors (Fig. 1D,F), suggesting that the ENU^+^ (NCpE) tumors have a more complex microenvironment.

To understand transcriptional changes imparted by ENU treatment of Nestin^+^ embryonic neural *Trp53*-deficient precursors, we performed a transcriptomic analysis comparing ENU^+^ vs. ENU^−^ tumors using bulk RNA sequencing. Gene set enrichment analysis revealed that immune-related pathways were significantly enriched in the ENU^+^ compared with ENU^−^ tumors (Supplementary Fig. 1F). To more precisely characterize the cellular composition of ENU⁺ tumors, we performed single-cell RNA sequencing (scRNA-seq) on four NCpE samples. Transcriptomic profiling at single-cell resolution revealed that the tumor samples had a large proportion of non-malignant immune cells, mirroring histological and bulk RNA-seq results (Fig. 1D, Supplementary Fig. 1G). The immune component comprised both lymphoid and myeloid cells, including tumor-associated microglia. The malignant population was dominated by cells that showed transcriptional programs reminiscent of cells of oligodendrocytic lineages, with smaller populations of cells demonstrating astrocytic and neuronal cell lineages (Supplementary Fig. 1H). Together, these data highlight that the NCpE model captures the diverse tumor microenvironmental milieu found in pediatric high-grade gliomas.

### Serial magnetic resonance imaging enables spatio-temporal tumor analysis

Live-animal magnetic resonance imaging (MRI) enables systematic, longitudinal tracking of tumor progression and allowed the detection of lesions at their earliest stages and their subsequent growth. We saw no lesions on MRI before 6 weeks, and called this period the premalignant stage. MRI analyses revealed that approximately 80% of NCpE tumors arose in supratentorial regions, while the remaining 20% originated in the infratentorial compartment [65] (Fig. 2A; Supplementary Fig. 2A–B), Survival outcomes did not differ significantly among mice with tumors originating in the cortex, striatum, or basally (Fig. 2B). However, mice with infratentorial tumors showed a modest but significant reduction in survival compared with those with cortical tumors, possibly a reflection of their earlier onset and faster progression, as well as of a potentially lower tolerance for tumor burden within this region.

**Figure 2.**
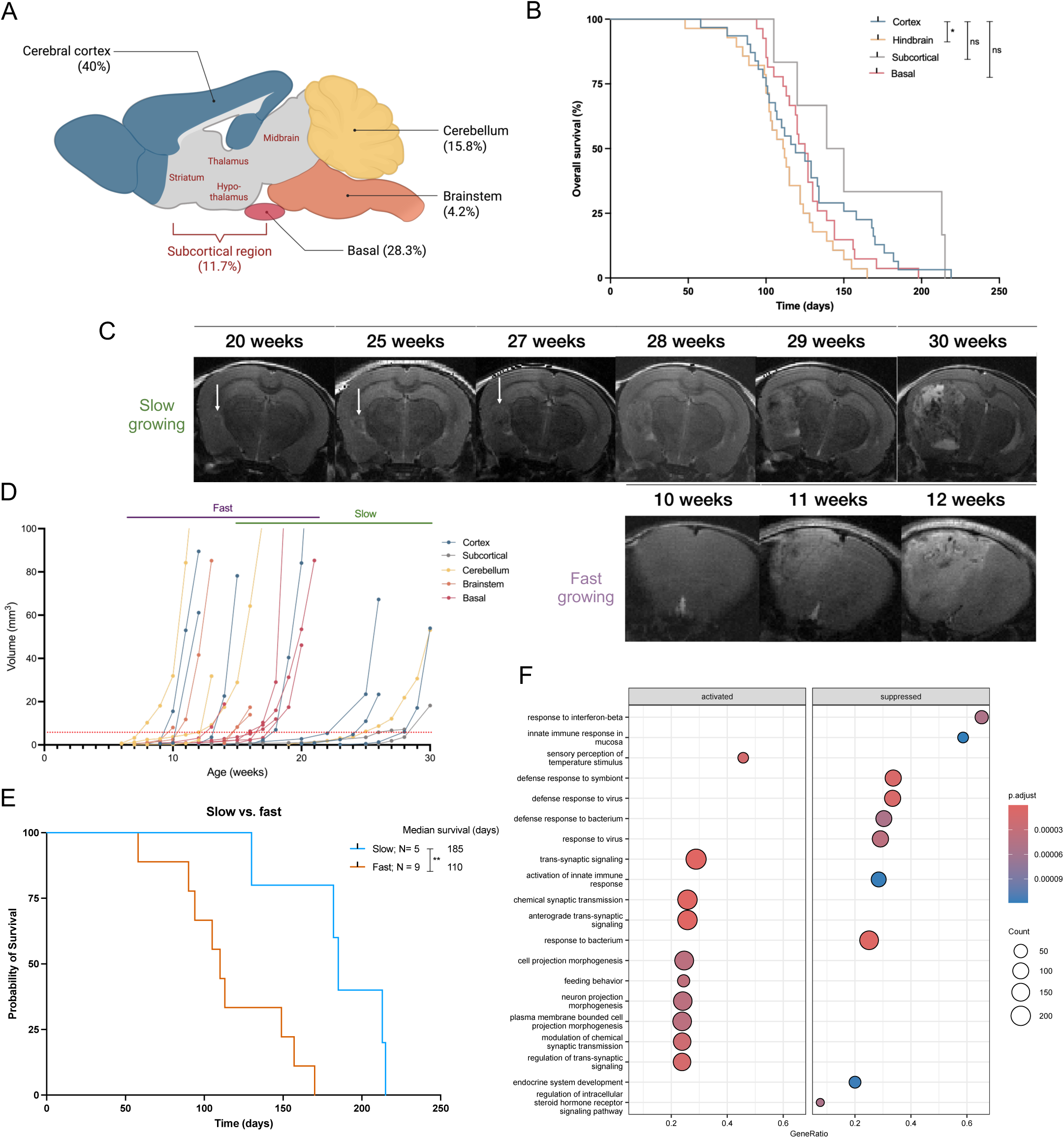
Characterization of NCpE tumor location and growth dynamics. (**A**) Anatomical distribution of NCpE end-stage tumors throughout the entire brain (total: n=120; cortex: n=48, cerebellum: 19, subcortical regions: n=14, brainstem: n=5, and trigeminal ganglion: n=34). Tumor incidence is depicted as a percentage of all identified tumors localized to the cerebral cortex. (**B**) Kaplan-Meier survival curves comparing NCpE tumor-bearing mice stratified by the anatomical location of the tumor (cortex: n=31, hindbrain: n=28, subcortical regions: n=6, and basal: n=27). Significance was estimated using the log-rank (Mantel-Cox) test. ns *p*≥0.05, \**p*<0.01. *p*=0.0198 (cortical vs. hindbrain tumors). (**C**) Longitudinal coronal MRI images of NCpE mice highlighting the progression of slow- and fast-growing tumors. Tumors are indicated by white arrows. (**D**) *In vivo* tumor growth kinetics between slow-(n=5) and fast-growing (n=6) NCpE tumors plotted as volume (mm^3^) versus age (weeks) and determined by serial MRI. (**E**) Kaplan-Meier survival curves comparing slow-(n=5) and fast-growing (n=9) NCpE tumors. Significance was estimated using the log-rank (Mantel-Cox) test. *p*=0.0044. (**F**) Gene ontology Biological Process (GO BP) enrichment analysis comparing representative enriched terms of highly expressed genes between slow- and fast-growing NCpE tumors. The dot size represents the number of genes and the color scale reflects the significance level.

We next performed differential gene expression analysis on bulk RNA sequenced samples to identify distinctly expressed markers between tumors arising in different brain regions. MRI demonstrated tumors that were confined to the cerebral cortex and others were found in the subcortical structures such as the thalamus and basal ganglia. The RNA expression supports that tumors retain a regional identity of expression defined by specific transcription factor expression. In the cortical and sub-cortical tumours, we observed expression of telencephalic markers such as *Foxg1* and *Lhx2,* with the expression of markers such as *Emx1*, *Emx2*, *Pax6*, and *Otx6* being more restricted to early lesions than end-stage tumors. The ventral forebrain marker *Nkx2-1* was more highly expressed in the sub-cortical tumors, and there was an absence of distinctly hindbrain markers such as *Hoxb2*, *Ptf1a*, *Atoh1*, and *Phox2b* in tumors of the cortex, sub-cortex and basal regions. (Supplementary Fig. 2C–D).

We also observed distinct temporal patterns of growth. All tumors had a long latent phase where size changed minimally week by week but then entered a short period of rapid growth that led to clinical symptoms. We studied a subset of lesions that appeared early (<12 weeks of age) and progressed rapidly from small lesions to end-stage tumors within 1–2 weeks. These occurred across different brain regions (N=14; 4 cortical, 6 hindbrain, 4 basal), and were classified as “fast” growing. We also studied a second, smaller subset that generally emerged later (>20 weeks of age) and required 3–4 weeks to reach end-stage size after initial detection; these were classified as “slow” growing, and occurred across regions (N=5; 3 cortical, 1 sub-cortical, 1 hindbrain) (Fig. 2C–D, Supplementary Fig. 2E). As expected, mice with fast-growing lesions had significantly shorter survival compared to those with slow-growing lesions (110 vs. 185 days, *p* = 0.0044; Fig. 2E. Interestingly, regardless of growth classification, all tumors exhibited a size threshold of approximately 5–8 mm^3^, beyond which rapid tumor expansion consistently occurred (Fig. 2D), revealing a critical stage in tumor growth that relates to a threshold tumor volume. A gene set enrichment analysis between genes that were differentially expressed in slow versus fast growing tumors at endpoint demonstrated an enrichment of innate and adaptive immune-related pathways in the fast-growing tumors compared to the slow, implicating the role of the tumor immune microenvironment in tumor growth (Fig. 2F). Taken together, the tumor growth dynamics encompass distinct patterns of proliferation that might model diverse tumor subtypes, with the fast-growing type being more enriched in inflammatory processes.

### End-stage tumors are transcriptomically and genomically more diverse than early-stage

Bulk RNA-sequencing of early and late-stage lesions revealed dynamic changes in transcriptional programs. Principal component analysis (PCA) separated tumors according to tumor progression stage, with early-stage lesions clustering close together, and end-stage tumors more widely distributed, reflecting their greater transcriptomic diversity (Fig. 3A–B).

**Figure 3.**
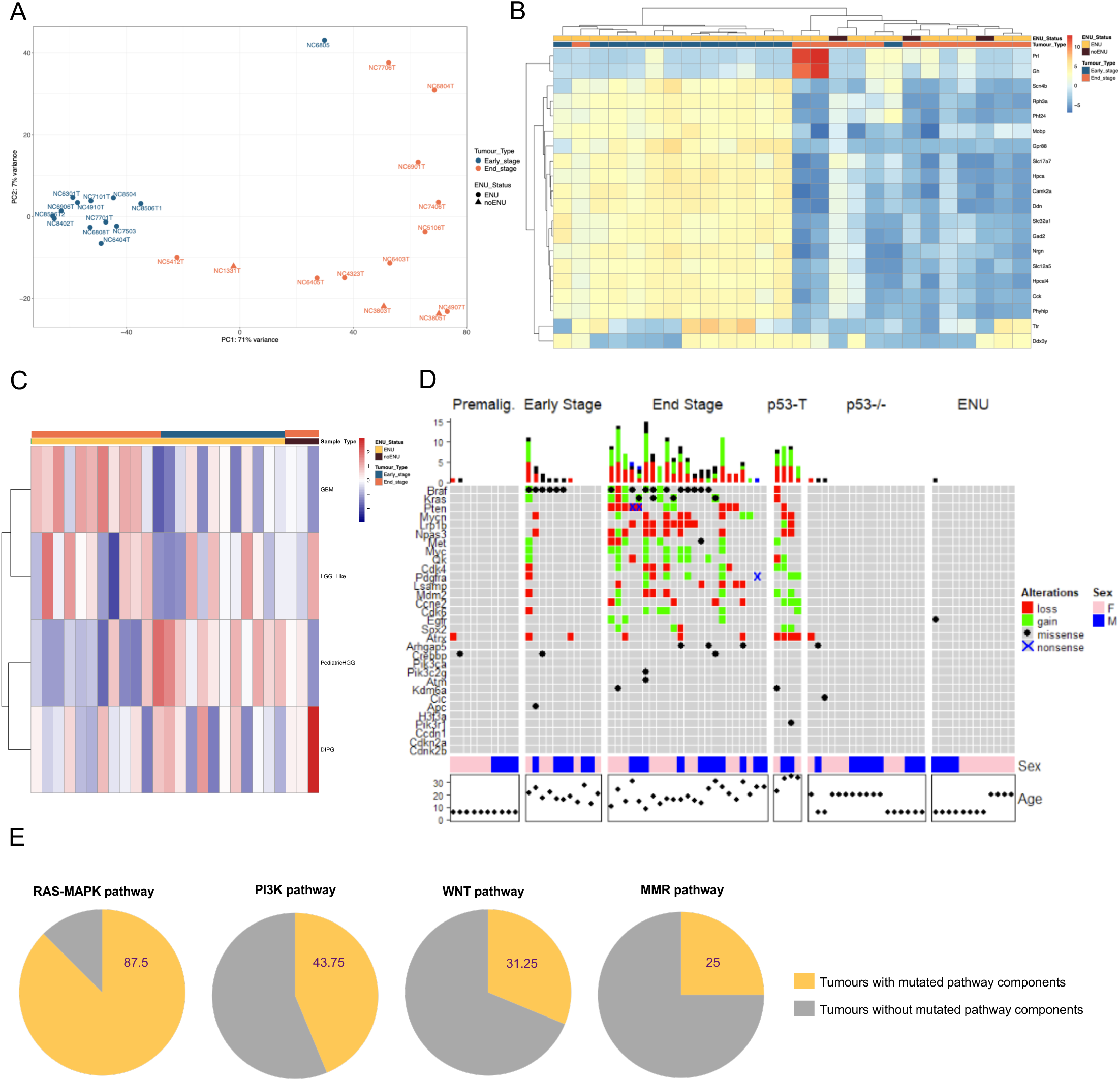
Transcriptomic and whole-exome sequence profiling of NCpE tumors reflects early and late-stage pediatric high-grade glioma biology. (**A**) Principal component analysis (PCA) of bulk RNA-seq data from NCp(E) tumors at early and end stage. Samples segregate primarily by tumor stage, with tight clustering in the early-stage tumors and more transcriptional variation in end-stage tumors. (**B**) Hierarchical heatmap showing the top 20 differentially expressed genes identified between early and late stage NCp tumors with and without ENU administration. Color scale indicates the scaled mean expression levels. (**C**) Gene set variation analysis (GSVA) of hallmark pathways in different brain cancer types: glioblastoma (GBM), low-grade glioma (LGG), pediatric high-grade glioma (pHGG), and diffused intrinsic pontine glioma (DIPG). Hierarchical heatmap depicts the normalized enrichment scores for each gene set across early-(n=11) and end-stage (n=12) NCpE tumors, as well as end-stage ENU^−^ (n=3) NCp tumors. (**D**) Oncoprint of genetic alterations (loss and gain of function, missense, and nonsense mutations) identified via whole-exome sequencing. Premalignant (n=10), early-stage (n=11), and end-stage (n=23) NCpE tumors, as well as NCp tumors (p53-T; n=4), p53 wildtype (p53-/-; n=17), and ENU only (n=12) controls were used in this analysis. Mutational burden in each tumor is summarized by the bar plot above. Sex and age are denoted in the annotation tracks below. (**E**) Summary of the proportion of NCpE tumors which harbor mutations in the: Ras-Mapk (87.5%), PI3K (43.75%), Wnt (31.25%) and MMR (25%) pathways.

Comparison of the mouse tumor data with human tumor datasets revealed a strong correlation between NCpE tumors and human high-grade gliomas, particularly those classified as H3- and IDH-wildtype (Fig. 3C and Supplementary Fig. 3A). End-stage tumors displayed transcriptomic features of adult glioblastoma (GBM), consistent with the known molecular overlap between pediatric and adult high-grade gliomas (Fig. 3C). Of the malignant cellular types previously identified in glioblastoma [65], we observed an enrichment in mesenchymal-like (MES1/2) and oligodendrocyte progenitor cell-like (OPC) in end-stage NCpE tumors, compared with neural progenitor cell-like (NPC) and astrocyte-like (AC) in early-stage lesions (Supplementary Fig. 3B), highlighting a more invasive, aggressive, and inflammatory phenotype as tumors progress from early to end stage. Interestingly, NCpE end-stage tumors had overlap with pediatric low-grade glioma (LGG) signatures, potentially highlighting the strong RAS-MAPK signaling activation in these lower grade tumors which is also seen in high grade tumors (Fig. 3C). Notably, early-stage lesions were also enriched for a pediatric high-grade glioma signature, suggesting that HGG drivers may emerge early in tumor evolution and may define potential clinically viable targets.

To determine the mutational profile imparted by ENU, we performed whole-exome sequencing (WES) of NCpE tumors. We hypothesized that this alkylating agent would produce monoallelic activating point mutations in drivers of GBM, in a *Trp53*-deficent context. We identified a high burden of single-nucleotide variants, predominantly C>T single nucleotide variants (SNVs), consistent with a common mutational signature of ENU (Supplementary Fig. 3C) [66]. Among glioma-related genes, we identified common mutations in *Braf*, *Kras*, *Nf1*, and *Nras*, suggesting MAPK pathway activation as a convergent oncogenic outcome (Fig. 3D), with almost 75% of ENU^+^ tumors having a mutated component in the RAS-MAPK signaling pathway (Fig. 3E).

Other mutated pathways include MycN, PI3K signaling, WNT signaling, and the DNA mismatch-repair (MMR) signaling pathways (Fig. 3E). Notably, activating *BrafV637E* mutations, orthologous to BRAFV600E in humans, were identified in a majority of the relatively mutationally quiet early-stage lesions (∼55%), and this phenomenon persisted into end-stage tumors (∼30%), suggesting a potential therapeutic vulnerability.

Genome-wide copy number variation (CNV) profiling revealed focal amplifications and deletions overlapping key glioma-associated regions, including amplification of *Egfr* and *Myc* (Fig. 3D). ENU^−^ tumors lacked point mutations, but exhibited copy number alterations including in *Sox2* and *Pdgfra*, which were also found in ENU^+^ tumors. Integration of SNV and CNV data highlighted a strong enrichment for RAS-MAPK pathway disturbance in the ENU^+^ lesions and tumors (Fig. 3D), highlighting the relevance of our model in recapitulating salient tumorigenic pathways.

### Transcriptomic and mutational convergence on the RAS-MAPK signaling pathway

Premalignant and early lesions displayed minimal copy number variation, whereas end-stage tumors acquired sub-clonal copy number alterations reminiscent of pediatric high-grade gliomas (Fig. 4A). Accordingly, CNV burden was increased in endpoint tumors, indicating that progressive chromosomal instability accompanies tumor evolution.

**Figure 4.**
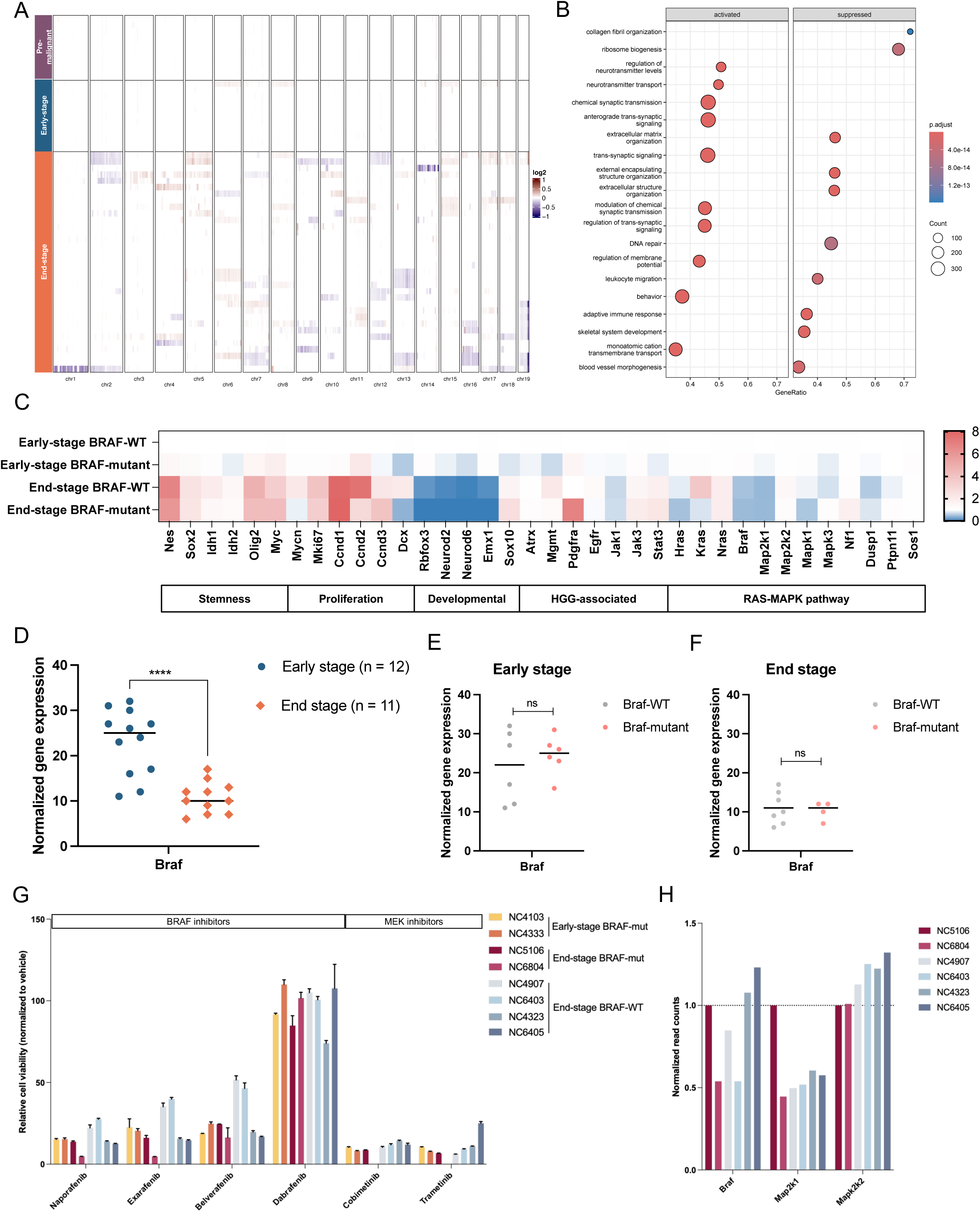
Early lesions are enriched for BRAF mutations and are susceptible to early intervention therapies targeting the Ras-Mapk pathway. (**A**) Heatmap showing copy number variations (CNV) of premalignant (n=1), early stage (n=11), and end stage (n=34) NCpE tumors in chromosomes 1-19. There is variability in copy number profiles across all tumors, with an enrichment in the end-stage tumors. (**B**) Gene ontology Biological Process (GO BP) enrichment analysis comparing representative enriched terms of highly expressed genes between BRAF-wildtype (WT) and BRAF-mutant NCpE tumors. The dot size represents the number of genes, and the color scale reflects the significance level. (**C**) Heatmap depicting the expression patterns of stemness, proliferation, developmental, HGG-associated, and Ras-Mapk-associated genes across early stage BRAF-WT (n=7), early stage BRAF-mutant (n=5), end stage BRAF-WT (n=7) and end stage BRAF-mutant (n=4) NCpE tumors. (**D–F**) Normalized expression levels of *Braf* across early and end-stage samples; **** *p* <0.0001. (**D**), between *Braf*-WT and *Braf* mutant samples in early stage samples (**E**), and between *Braf*-WT and *Braf* mutant samples in end stage samples (**F**). (**G**) Relative cell viability of NCpE-derived tumor cells and to BRAF inhibitors: Naporafenib, Exarafenib, Belverafenib, and Dabrafenib; and MEK inhibitors: Cobimetinib and Trametinib (all used at 5 mM). NCpE-derived cells were of early-stage BRAF-mutant (N=2), end-stage BRAF-mutant (N=2), and end-stage BRAF-WT (N=4) status. Data shown represent mean ± [SEM] normalized to the DMSO-treated control. (**H**) Normalized expression levels of *Braf*, *Map2k1*, and *Map2k2* across the end stage NCpE-tumor derived cell lines.

Comparing early-stage and end-stage tumors transcriptomically, gene ontology and gene set enrichment analyses showed enrichment of DNA repair, adaptive immune response, ribosome biogenesis, and blood vessel morphogenesis, consistent with hallmark cancer programs becoming activated during progression, while early lesions retained developmental and neural progenitor signatures (Fig. 4B). At the gene level, stemness-associated factors such as *Sox2*, *Idh1*, *Idh2*, and *Nestin*, as well as the proliferation markers *Ccnd1*, *Ccnd2*, *Mki67*, were upregulated, whereas the markers of neuronal specification *Neurod1*, *Neurod6, Rbfox3* were downregulated (Fig. 4C). Notably, early lesions exhibited significantly higher *Braf* expression compared with end-stage. This pattern was independent of *Braf* mutation status, and stratification by *Braf* mutation did not significantly impact the median survival of tumor-bearing mice at end-stage (Supplementary Fig. 4A–B), further supporting the notion that RAS-MAPK signaling is an early driver in these tumors (Fig. 4D–F). Indeed, *BRAF* mutations are frequently found in pediatric low-grade gliomas, with *BRAF*-mutant LGGs more likely to progress to high-grade gliomas with cooperating mutations [67].

Collectively, these results demonstrate that the NCpE model reveals interesting features at different stages of gliomagenesis, stemming from a random mutagenesis approach in a *Trp53*-null context. Tumors resemble many key aspects of both adult and pediatric GBMs from a transcriptomic and mutational perspective. The recurrent alterations identified in RAS-MAPK pathway from this mutational approach lend strong credence to activation of this pathway, in neural precursors, as a key bottle neck in driving glioma.

### NCpE tumor cells respond to BRAF inhibitors

Given the higher *Braf* expression observed in early-stage lesions compared with end-stage tumors, we compared the responsiveness of primary cells derived from both early and end-stage lesions (N=3 and N=6, respectively), encompassing a variety of different genotypes. We studied four BRAF inhibitors: Naporafenib, Exarafenib, Belvarafenib, and Dabrafenib (Supplementary Fig. 4C) (OICR), and two commercially available MEK inhibitors, Cobimetinib and Trametinib. Notably, Dabrafenib is clinically used to treat BRAF-mutant pediatric high-grade gliomas, although resistance frequently emerges in tumors that become malignant [68]. *Braf* expression was significantly higher in early lesions compared with end-stage tumors (Fig. 4D–F), and cells derived from early-stage *Braf*-mutant lesions displayed sensitivity to both BRAF and MEK inhibition. Among end-stage lines, BRAF-mutant cells were more sensitive than lines that did not carry the BRAF mutation (Fig. 4G). In all cases, the best predictor of responsiveness to BRAF and MEK inhibition, was the level of *Braf* and *Map2k1/2* (homologous to human *MEK1/2*) expression respectively (Fig. 4H).

To assess potential clinical relevance of these findings, we also tested BRAF inhibition in high-grade glioma lines derived from pediatric primary human tumors (N=4). These lines exhibited molecular features of pHGG, including chromosome 10 loss (G477), chromosome 7 gain (G626), and missense mutations in BRAF (G626), GRIN2A, WNT2, and PRDM9. Upon treatment, they showed variable response that also correlated with their *BRAF* expression pattern (Supplementary Fig. 4D–E). Interestingly, these findings show that HGG lines derived from our NCpE model display BRAF-inhibitor sensitivity that correlates with *Braf* expression rather than strictly with mutational status. Moreover, consistent with clinical experience, BRAF inhibition was more effective in early lesions, when tumors had higher BRAF expression. Together, our model effectively captures stages of gliomagenesis and reveals potential windows of vulnerability with enhanced therapeutic potential.

## Discussion

High-grade gliomas (HGGs) remain among the most aggressive human cancers, with poor prognoses despite advances in molecular classification and targeted therapies. One reason for continued failure of treatment is that we still lack understanding of how these tumors initiate and progress, and what the true molecular drivers of disease are. By the time HGGs present clinically, they are highly genetically and cellularly diverse, with multiple genetic clones mixed together, that driver cell types and their clones are obscured. Human cell models approximating endpoint disease can be studied in xenograft assays, and are accessible for various drug screens, but they do not provide mechanistic insight into how normal cells transform into malignant populations, nor how preneoplastic states evolve to malignant clones. In short, the early steps of gliomagenesis cannot be readily accessed in human subjects.

In mice, tumors are engineered typically by simultaneous co-expression or loss of function of specific HGG-associated genes, but it is unlikely that the human disease results from an abrupt simultaneous loss of multiple key genes in normal cells, and disease likely follows a slower evolutionary process. High-grade gliomas likely arise through the sequential acquisition of intrinsic alterations in genomic sequence coupled with extrinsic influences that exert themselves over a long period, and the cell context in which these changes occur and evolve is likely critical.

We therefore generated an autocthonous model where genetic mutations are more randomly acquired and where we could study multiple stages of the disease, in an immunocompetent context. We used a *Nestin^Cre/+^*;*Trp53^fl/fl^* mouse model layered with embryonic ENU mutagenesis. This combined approach mirrors aspects of gliomagenesis that are obscure in other more typical GEMMs.

This model reliably produces brain tumors that recapitulate the histopathological and molecular features of human HGGs. Following a latency period where no tumor could be seen on imaging, all mice develop tumors within a reproducible timeframe facilitating genotypic and phenotypic analysis through stages of tumor development. ENU exposure dramatically accelerates tumor onset, increases penetrance, and generates more immune cell-rich tumors, providing a framework to study how stochastic mutagenic events intersect with genetic predisposition to initiate gliomagenesis. Critically, by preserving the native brain microenvironment and immune system, elements often absent from xenograft or transgenic models, this system provides an immunocompetent and developmentally relevant context in which to study tumor evolution.

Integration of live MR imaging further enables non-invasive monitoring of tumor emergence and progression, and the capturing of samples at premalignant, early-, and end-stage phases for downstream analyses. Early growth dynamics in HGGs are rarely observed in human studies [69], and tumors are often several centimeters in diameter by the time of diagnosis. In contrast, our approach permits observation of lesions as small as 0.3 mm in size, potentially revealing critical windows of vulnerability during early tumor developmental stages.

Whole-exome sequencing revealed that the mutational landscape evolves progressively across tumor stages, with the frequency of copy number alterations increasing from premalignant to end-stage lesions. The activation of mutations in genes of the RAS–MAPK pathway in the ENU^+^ samples, in which disease progression is accelerated, reinforces how critical activation of this pathway is for gliomagenesis. In particular, activating *BrafV637E* mutations, orthologous to human BRAFV600E, emerged early and persisted into end-stage tumors, re-enforcing how important activation this oncogene is for gliomagenesis, particularly in children. Additional oncogenic mutations and copy number alterations arose later, establishing a temporal hierarchy of genetic events and reflecting the genomic instability and complexity in later-stage tumors that may underlie recalcitrance to therapy. It is well known that high-grade gliomas at clinical presentation are characterized by extensive copy number alterations, and this model may allow discernment of the mechanisms by which tumors evolve related to the acquisition of this dramatic genomic complexity which seems coincident with explosive tumor growth.

Transcriptomic profiling revealed a clear separation between early and end-stage lesions, highlighting transcriptional differences between tumor developmental stages. Pathway analyses demonstrated persistent activation of MAPK target genes, and increasing activation of proliferation-associated and stemness programs during tumor progression, while markers of neuronal specification were downregulated, reflecting corruption of the developmental hierarchy and suppression of differentiation amongst malignant cells during tumorigenesis.

The cooperation between TRP53 loss and MAPK pathway activation in this model recapitulates central pathogenic mechanisms observed in both pediatric and adult gliomas. Our findings suggest that MAPK pathway activation is an early, targetable event in gliomagenesis, and raises the possibility that targeted pharmacologic inhibition earlier in tumorigenic development could mitigate progression to high-grade disease.

Future directions will focus on increasing temporal resolution through additional intermediate lesion sampling and expanded longitudinal MRI datasets to define early growth kinetics more precisely. Single-cell and spatial transcriptomic approaches will help disentangle tumor-microenvironment interactions. Investigating the impact of ENU on non-tumor compartments, including microglia and astrocytes, may reveal whether mutations also indirectly promote tumorigenesis through micro-environmental remodeling. Finally, the *in vitro* results indicating that BRAF inhibition is more effective in early-stage tumors compared with end-stage will be tested *in vivo*, potentially in combination with other viable targets.

In summary, the ENU–p53 gliomagenesis model provides a robust, reproducible, and immunocompetent platform to study glioma evolution. It recapitulates key genetic and histopathological hallmarks of human disease, allows real-time visualization of early-to-late lesions, and converges on critical patient-relevant mutations and tumor signaling pathways. The combination of internal littermate controls and reproducible tumor onset establishes an internally controlled system to interrogate how random mutational events generated by an external agent interact with genetic susceptibility. Collectively, these features position this model as a highly tractable platform for investigating early tumorigenic mechanisms, identifying therapeutic vulnerabilities, and testing strategies to intercept gliomagenesis prior to full-blown malignancy, with RAS-MAPK pathway activation emerging as a particularly compelling target for early intervention.

## Acknowledgements

We would like to thank the SickKids Imaging Facility, SickKids Laboratory Animal Services, and the UHN Spatio-Temporal Targeting and Amplification of Radiation Response (STTARR) Facility for contributions to this work. We would also like to thank Dr. Warren Foltz at the STTARR facility for acquiring the mouse magnetic resonance images (MRI) and advice on MRI analysis. We thank present and former members of the Dirks lab for all their support, including Vera Luo for assistance with tissue culture. Research was supported but Stand Up To Cancer (SU2C) Canada Cancer Stem Cell Dream Team Research Funding (SU2C-AACR-DT-19-15) provided by the Government of Canada through Genome Canada and the Canadian Institute of Health Research and Stand up to Cancer Pediatric Cancer Catalyst grant. K.D. is supported by the Garry Hurvitz Centre for Brain and Mental Health (GH-CBMH) Brain and Mental Health Outcomes Catalyst Grant.

