## Supplemental figures for "Convergence on BRAF and MAPK Signaling in Glioma Development in a P53-ENU model"

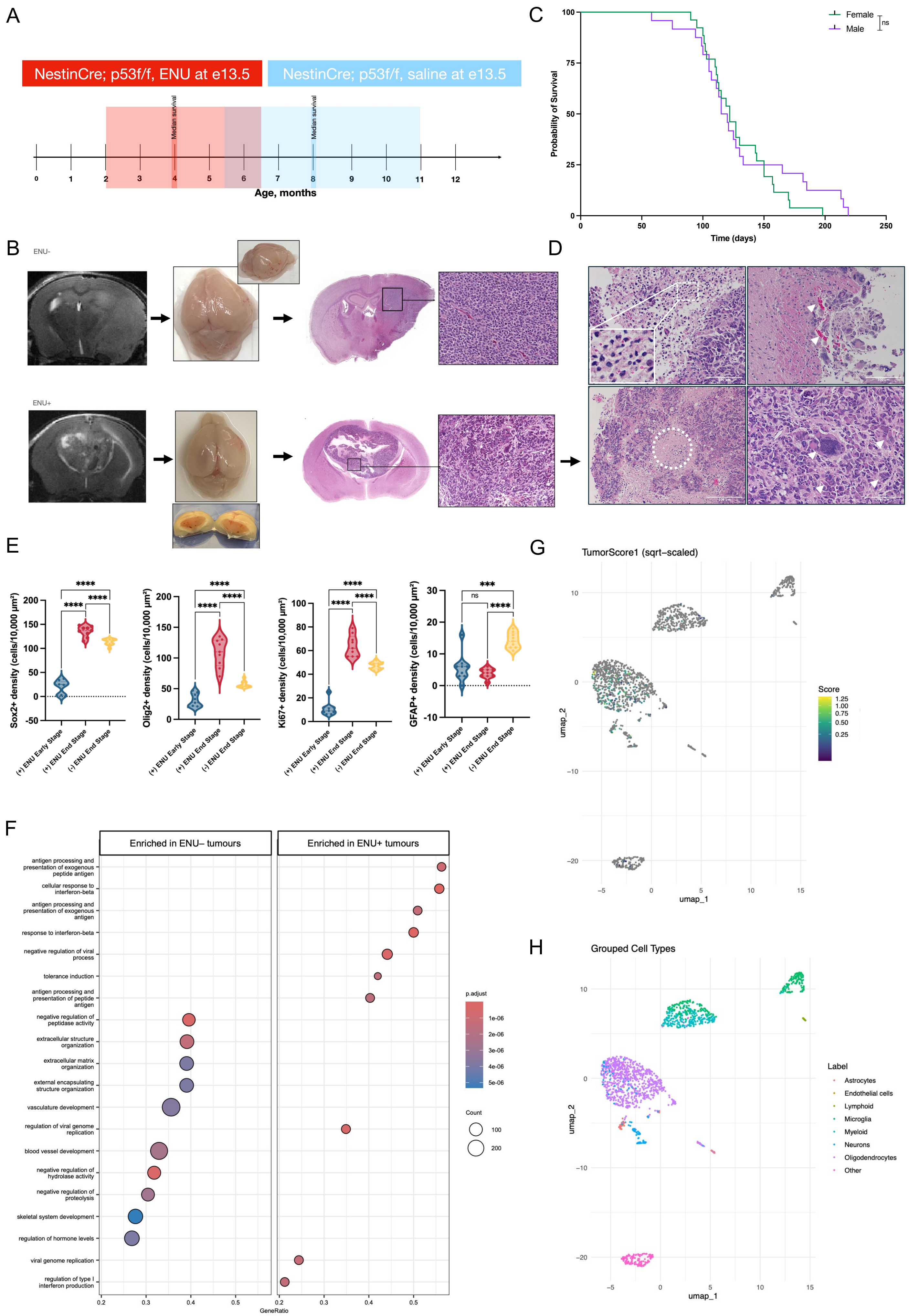

A
NestinCre; p53f/f, ENU at e13.5
NestinCre; p53f/f, saline at e13.5
Median survival
Median survival
Age, months
C
Female
Male
ns
Probability of Survival
Time (days)
B
ENU-
ENU+
D
E
Sox2+ density (cells/10,000 μm²)
Olig2+ density (cells/10,000 μm²)
Ki67+ density (cells/10,000 μm²)
GFAP+ density (cells/10,000 μm²)
(+) ENU Early Stage
(+) ENU End Stage
(-) ENU End Stage
F
Enriched in ENU– tumours
Enriched in ENU+ tumours
antigen processing and presentation of exogenous peptide antigen
cellular response to interferon-beta
antigen processing and presentation of exogenous antigen
response to interferon-beta
negative regulation of viral process
tolerance induction
antigen processing and presentation of peptide antigen
negative regulation of peptidase activity
extracellular structure organization
extracellular matrix organization
external encapsulating structure organization
vasculature development
regulation of viral genome replication
blood vessel development
negative regulation of hydrolase activity
negative regulation of proteolysis
skeletal system development
regulation of hormone levels
viral genome replication
regulation of type I interferon production
GeneRatio
p.adjust
Count
G
TumorScore1 (sqrt–scaled)
Score
umap_1
umap_2
H
Grouped Cell Types
Label
Astrocytes
Endothelial cells
Lymphoid
Microglia
Myeloid
Neurons
Oligodendrocytes
Other

Supplementary Figure 1

(**A**) Graphic depicting the window of the age of onset and median survival of *Nestin*$^{Cre/+}$;*Trp53*$^{fl/fl}$ with ENU (red) or saline (blue) administered intraperitoneally on embryonic day 13.5 (NCpE). (**B**) Representative whole brain tissue sections collected at end stages from NCp and NCpE mice, photographed, and stained for H&E. H&E scale bars: 1000 µm. (**C**) Kaplan-Meier survival curve of male and female NCpE mice (females: n=26, males: n=23). Significance was estimated using the log-rank (Mantel-Cox) test. Chi square =0.4458, *p*=0.5043 (non-significant). (**D**) Representative H&E images from NCpE tumours. Dotted white lines represent characteristic features of high-grade gliomas: hypercellularity, nuclear atypia, necrosis, microvascular proliferation; H&E scale bars: 1000 µm. (**E**) Quantification of various markers (Sox2, Olig2, Ki67, and GFAP) present in the early-stage lesion or end-stage tumor core. For the quantification, three 100 µm x 100 µm regions were evaluated from three independent biological samples per group. Dotted lines within the violin plots denote the median. Two-tailed unpaired *t-test*; ****p*<0.001, *****p*<0.0001. (**F**) Gene ontology Biological Process (GO BP) enrichment analysis comparing representative enriched terms of highly expressed genes between ENU$^{-}$ (NCp) and ENU$^{+}$ (NCpE) tumors. The dot size represents the number of genes and the color scale reflects the significance level. (**E–H**) scRNAseq analysis on four NCpE mouse tumours; unsupervised clustering using Uniform Manifold Approximation and Projection (UMAP) performed using tumor scoring metric identifying malignant cell populations (E) as well as clustering of cell types in dataset (F).

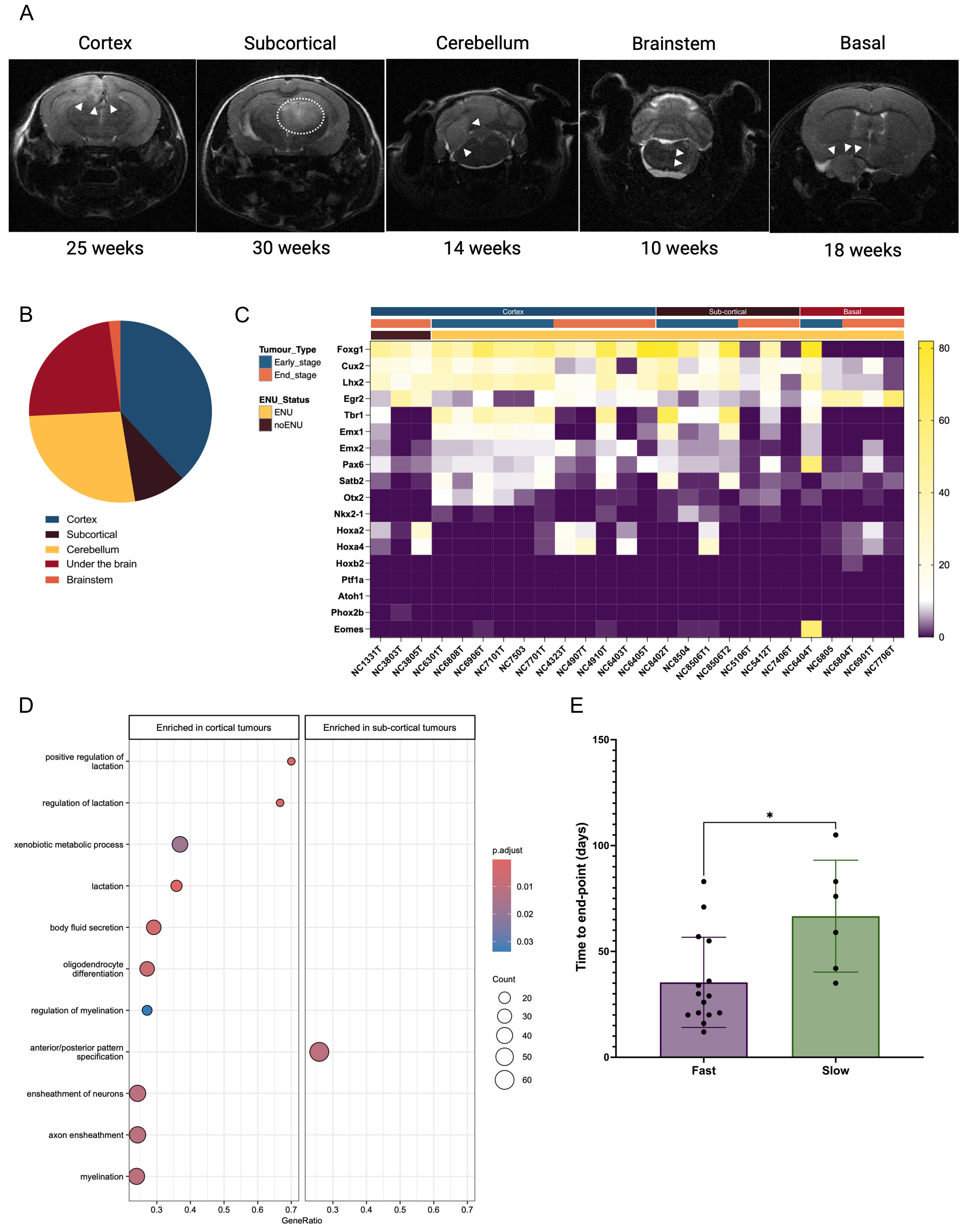
A
Cortex
Subcortical
Cerebellum
Brainstem
Basal
25 weeks
30 weeks
14 weeks
10 weeks
18 weeks
B
Cortex
Subcortical
Cerebellum
Under the brain
Brainstem
C
Tumour_Type
Early_stage
End_stage
ENU_Status
ENU
noENU
Cortex
Sub-cortical
Basal
Foxg1
Cux2
Lhx2
Egr2
Tbr1
Emx1
Emx2
Pax6
Satb2
Otx2
Nkx2-1
Hoxa2
Hoxa4
Hoxb2
Ptf1a
Atoh1
Phox2b
Eomes
NC1331T
NC3803T
NC3805T
NC6301T
NC6808T
NC6906T
NC7101T
NC7503
NC7701T
NC4323T
NC4907T
NC4910T
NC6403T
NC6405T
NC8402T
NC8504
NC8506T1
NC8506T2
NC5106T
NC5412T
NC7406T
NC6404T
NC6805
NC6804T
NC6901T
NC7706T
80
60
40
20
0
D
Enriched in cortical tumours
Enriched in sub-cortical tumours
positive regulation of lactation
regulation of lactation
xenobiotic metabolic process
lactation
body fluid secretion
oligodendrocyte differentiation
regulation of myelination
anterior/posterior pattern specification
ensheathment of neurons
axon ensheathment
myelination
0.3
0.4
0.5
0.6
0.7
GeneRatio
p.adjust
0.01
0.02
0.03
Count
20
30
40
50
60
E
Time to end-point (days)
150
100
50
0
*
Fast
Slow

Supplementary Figure 2

(**A**) Representative longitudinal coronal MRI images of NCpE mice illustrating the regional distribution of tumors. Tumors are indicated by white arrows. (**B**) Pie chart showing the distribution of tumor anatomical location (cortex: n=31, hindbrain: n=28, subcortical regions: n=6, and basal: n=27). (**C**) Heatmap depicting the expression patterns of cortical, sub-cortical and hindbrain markers across early stage NCpE and end stage NCp and NCpE tumors distributed across the cortex, sub-cortex, and basal regions. (**D**) Gene ontology Biological Process (GO BP) enrichment analysis comparing representative enriched terms of highly expressed genes between cortical and sub-cortical originating $ENU^{+}$ (NCpE) tumors. The dot size represents the number of genes and the color scale reflects the significance level. (**E**) Average time from lesion initiation to end-stage in fast- versus slow-growing NCpE tumour groups. Time was defined as the number of days between first detection of an early lesion by MRI and the final end-stage MRI (tumour volume capped at 100 $mm^3$), calculated exclusively. The fast-growing group included n=15 tumours (n=4 cortex; n=4 brainstem; n=3 cerebellum; n=4 basal), and the slow-growing group included n=6 tumours (n=3 cortex; n=1 cerebellum; n=2 subcortical). Mean time to end-stage was 35.4 days for fast-growing tumours and 66.7 days for slow-growing tumours. Statistical significance was determined using Welch's t-test ($p$=0.0334).

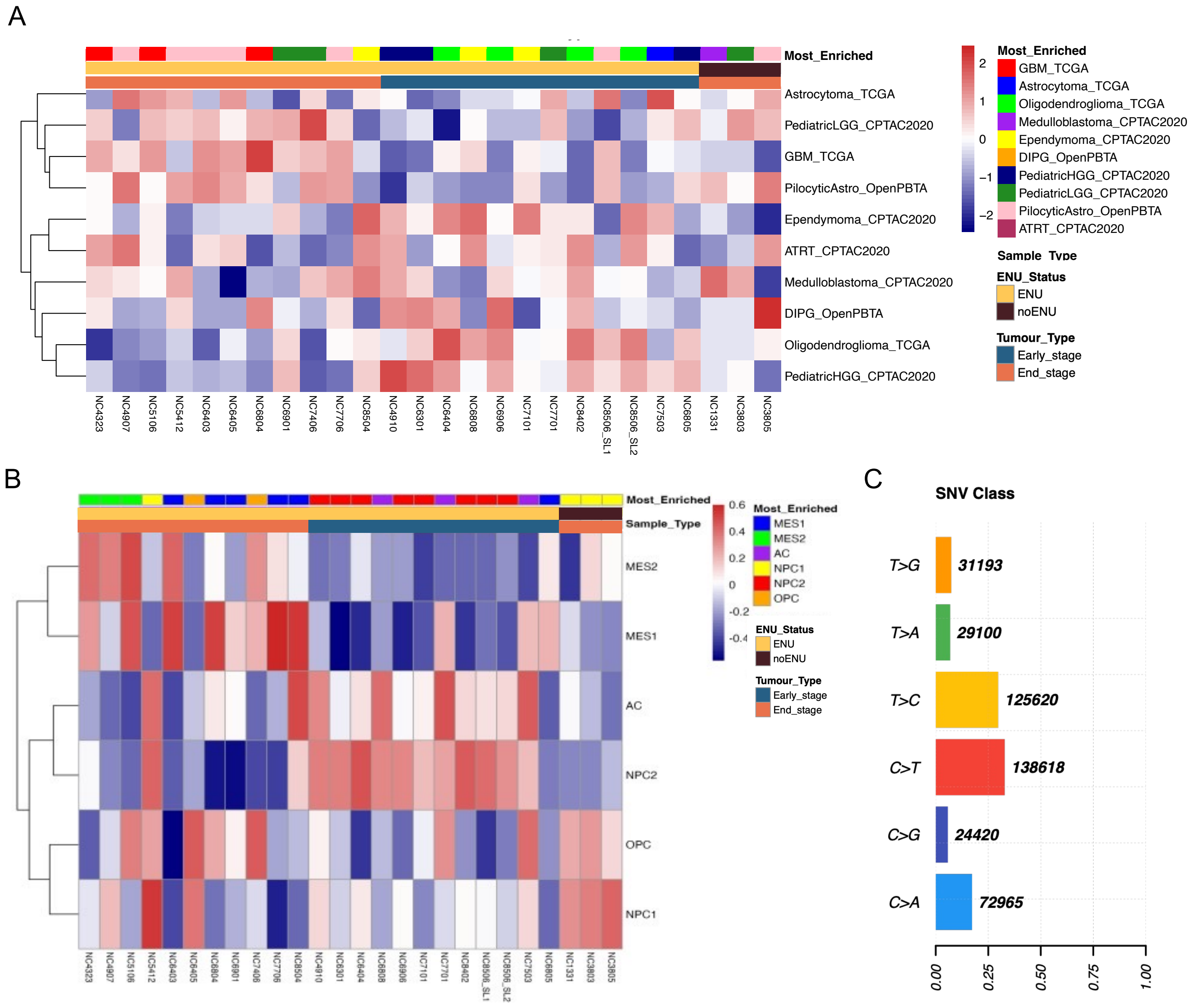
A
Most_Enriched
Astrocytoma_TCGA
PediatricLGG_CPTAC2020
GBM_TCGA
PilocyticAstro_OpenPBTA
Ependymoma_CPTAC2020
ATRT_CPTAC2020
Medulloblastoma_CPTAC2020
DIPG_OpenPBTA
Oligodendroglioma_TCGA
PediatricHGG_CPTAC2020
NC4323
NC4907
NC5106
NC5412
NC6403
NC6405
NC6804
NC6901
NC7406
NC7706
NC8504
NC4910
NC6301
NC6404
NC6808
NC6906
NC7101
NC7701
NC8402
NC8506_SL1
NC8506_SL2
NC7503
NC6805
NC1331
NC3803
NC3805
2
1
0
−1
−2
Most_Enriched
GBM_TCGA
Astrocytoma_TCGA
Oligodendroglioma_TCGA
Medulloblastoma_CPTAC2020
Ependymoma_CPTAC2020
DIPG_OpenPBTA
PediatricHGG_CPTAC2020
PediatricLGG_CPTAC2020
PilocyticAstro_OpenPBTA
ATRT_CPTAC2020
Sample_Type
ENU_Status
ENU
noENU
Tumour_Type
Early_stage
End_stage
B
Most_Enriched
Sample_Type
MES2
MES1
AC
NPC2
OPC
NPC1
0.6
0.4
0.2
0
-0.2
-0.4
Most_Enriched
MES1
MES2
AC
NPC1
NPC2
OPC
ENU_Status
ENU
noENU
Tumour_Type
Early_stage
End_stage
C
SNV Class
T>G
31193
T>A
29100
T>C
125620
C>T
138618
C>G
24420
C>A
72965
0.00
0.25
0.50
0.75
1.00

Supplementary Figure 3

(**A–B**) Gene set variation analysis (GSVA) of hallmark pathways in different brain cancer types (A) and the transcriptional cellular states reported found in glioblastoma (GBM) (Neftel *et al,* 2022) (**B**). AC: astrocyte-like, MES1: mesenchymal-1-like, MES2: mesenchymal-2-like, OPC: oligodendrocyte-progenitor-like, NPC1: neural-progenitor-1-like, and NPC2: neural-progenitor-2-like. Hierarchical heatmap depicts the normalized enrichment scores for each gene set across early- (n=11) and end-stage (n=12) NCpE tumors, as well as end-stage ENU– (n=3) NCp tumors. (**C**) Relative contribution of the six SNV classes to the tumor mutational burden found in the early and late-stage NCpE tumors through whole-exome sequencing (WES) analysis. Somatic SNV classes are calculated based on the substitution type. Colors represent the type of substitution.

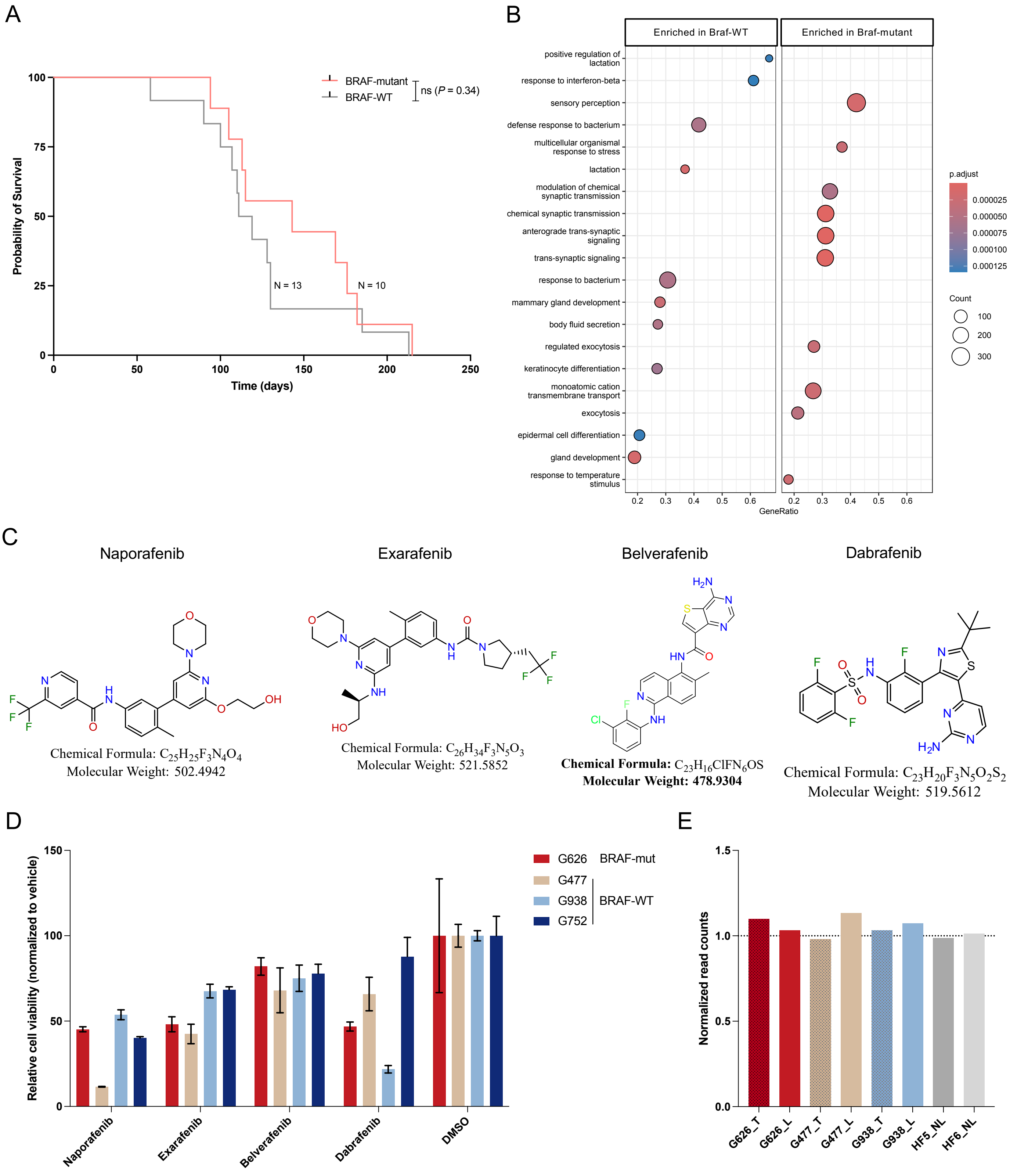
A
Probability of Survival
Time (days)
BRAF-mutant
BRAF-WT
ns (P = 0.34)
N = 13
N = 10
B
Enriched in Braf-WT
Enriched in Braf-mutant
positive regulation of lactation
response to interferon-beta
sensory perception
defense response to bacterium
multicellular organismal response to stress
lactation
modulation of chemical synaptic transmission
chemical synaptic transmission
anterograde trans-synaptic signaling
trans-synaptic signaling
response to bacterium
mammary gland development
body fluid secretion
regulated exocytosis
keratinocyte differentiation
monoatomic cation transmembrane transport
exocytosis
epidermal cell differentiation
gland development
response to temperature stimulus
GeneRatio
p.adjust
Count
C
Naporafenib
Chemical Formula: $C_{25}H_{25}F_3N_4O_4$
Molecular Weight: 502.4942
Exarafenib
Chemical Formula: $C_{26}H_{34}F_3N_5O_3$
Molecular Weight: 521.5852
Belverafenib
Chemical Formula: $C_{23}H_{16}ClFN_6OS$
Molecular Weight: 478.9304
Dabrafenib
Chemical Formula: $C_{23}H_{20}F_3N_5O_2S_2$
Molecular Weight: 519.5612
D
Relative cell viability (normalized to vehicle)
G626 BRAF-mut
G477
G938 BRAF-WT
G752
Naporafenib
Exarafenib
Belverafenib
Dabrafenib
DMSO
E
Normalized read counts
G626_T
G626_L
G477_T
G477_L
G938_T
G938_L
HF5_NL
HF6_NL

Supplementary Figure 4

(**A**) Kaplan-Meier survival curve of *Braf*-mutant and *Braf* wild-type (WT) NCpE mice (*Braf*-mutant: n=10, *Braf*-WT: n=13). Significance was estimated using the log-rank (Mantel-Cox) test. Chi square =0.9099, $p$=0.3401 (non-significant). (**B**) Gene ontology Biological Process (GO BP) enrichment analysis comparing representative enriched terms of highly expressed genes between *Braf*-mutant and *Braf* wild-type $ENU^{+}$ (NCpE) tumors. The dot size represents the number of genes and the color scale reflects the significance level. (**C**) Chemical structures, chemical formula, and molecular weights of the BRAF inhibitors obtained from the Ontario Institute of Cancer Research: Naporafenib, Exarafenib, Belverafenib, and Dabrafenib. (**D**) Relative cell viability of human pediatric high-grade glioma (pHGG) cell lines to BRAF inhibitors: Naporafenib, Exarafenib, Belverafenib, and Dabrafenib; and MEK inhibitors: Cobimetinib and Trametinib (all used at 5 mM). Human pHGG cells were of BRAF-WT (N=3) and BRAF-mutant (N=1, G626) status. Data shown represent mean ± [SEM] normalized to the DMSO-treated control. (**E**) Normalized expression levels *Braf* across the human pHGG primary tumor tissue and derived cell lines.
